# Concurrent feature-specific reactivation within the hippocampus and neocortex facilitates episodic memory retrieval

**DOI:** 10.1101/2021.02.28.433226

**Authors:** Michael B. Bone, Bradley R. Buchsbaum

## Abstract

The hippocampus is a key brain region for the storage and retrieval of episodic memories, but how it performs this function is unresolved. According to the hippocampal indexing theory, the hippocampus stores an event-specific index of the pattern of neocortical activity that occurred during perception. During retrieval, reactivation of the index by a partial cue facilitates the reactivation of the associated neocortical pattern. Therefore, event-specific retrieval requires joint reactivation of the hippocampal index and the associated neocortical networks. To test this theory, we examine the relation between performance on a recognition memory task requiring retrieval of image-specific visual details and feature-specific reactivation within the hippocampus and neocortex. We show that trial-by-trial recognition accuracy correlates with neural reactivation of low-level features (e.g. luminosity and edges) within the posterior hippocampus and early visual cortex for participants with high recognition lure accuracy. As predicted, the two regions interact, such that recognition accuracy correlates with hippocampal reactivation only when reactivation co-occurs within the early visual cortex (and vice-versa). In addition to supporting the hippocampal indexing theory, our findings show large individual differences in the features underlying visual memory and suggest that the anterior and posterior hippocampus represents gist-like and detailed features, respectively.

---

The ability to mentally reexperience vivid imagery from a past event, a defining feature of episodic memory, has been associated with the reactivation of neural activity that occurred during the original episode. Neocortical reactivation has been shown to represent the content of episodic memory (Buchsbaum et al., 2012; Kuhl, Bainbridge, & Chun, 2012; Johnson & Johnson, 2014; St-Laurent et al., 2014; Cabeza, Ritchey & Wing, 2015; Bone et al., 2019), including low-level visual properties such as edge orientation and luminosity (Harrison & Tong, 2009; Albers et al., 2013; Naselaris et al., 2015; Bone, Ahmad & Buchsbaum, 2020), as well as high-level semantic properties (Reddy, Tsuchiya & Serre, 2010; Cichy, Heinzle & Haynes, 2011; Bone, Ahmad & Buchsbaum, 2020). Consistent with the role of the hippocampus in memory storage, reactivation has also been found within the hippocampus (Chadwick, Hassabis, Weiskopf & Maguire, 2010). Studies have shown hippocampal reactivation representing specific images (Mack & Preston, 2016), event-specific associations (Tompary, Duncan & Davachi, 2016), object category (Liang & Preston, 2017), context (Ritchey, Montchal, Yonelinas & Ranganath, 2015), location (Brown et al., 2016), and event details (Thakral, Madore, Addis, & Schacter, 2019).

Leading accounts of hippocampal function posit that the region stores an event-specific index of the pattern of neocortical activity that occurred during perception, thereby facilitating the encoding of arbitrary associations between an event’s constituent features that can later be retrieved by the reactivation of a subset of the encoded features (Teyler & DiScenna, 1986, Nadel, Samsonovich, Ryan & Moscovitch, 2000; Sekeres, Winocur & Moscovitch, 2018). To prevent interference from similar memories with shared features, overlap between indexes is minimized by ‘pattern separation’ processes primarily carried out by the dentate gyrus and CA3 hippocampal subfields (Bakker, Kirwan, Miller & Stark, 2008; Rolls, 2013). The patterns that can be indexed and the features that they represent are determined by the physical connections between the hippocampus and neocortex. This connectivity varies along the longitudinal axis, with the posterior hippocampus (pHC) reciprocally linked to sensory regions of the posterior neocortex, and the anterior hippocampus (aHC) connected to anterior neocortical structures including the anterior lateral temporal cortex which mediates semantic information, and the ventromedial prefrontal cortex which is implicated in the representation of schemas (Poppenk & Moscovitch, 2011; Kier, Staib, Davis & Bronen, 2004; Catenoix, Magnin, Mauguiere, & Ryvlin, 2011; Poppenk, Evensmoen, Moscovitch, & Nadel, 2013). Based on differences in connectivity and receptive field size (Kjelstrup et al., 2008) along the hippocampal longitudinal axis, researchers (McCormick et al., 2013; Poppenk, Evensmoen, Moscovitch, & Nadel, 2013) postulate that the pHC indexes the fine-grained perceptual features of an event which constitute vivid, perceptually rich memories, whereas the aHC indexes coarse-grained features, which support gist-like memories. Although the bulk of experimental evidence supports this hypothesis (Schlichting, Mumford, & Preston, 2015; Evensmoen et al. 2015; Brunec et al., 2018; Sekeres, Winocur & Moscovitch, 2018; Grady, 2019), at least one recent finding implicates the aHC and pHC in the representation of detailed and gist-like memories, respectively (Dandolo & Schwabe, 2018).

Here we examine the predictions of indexing theory and the contributions of the aHC and pHC to detailed visual memory. We combined functional magnetic resonance imaging (fMRI) and measures of neural reactivation applied to a challenging recall and recognition task. We defined reactivation in two ways: image-specific and feature-specific reactivation. Image-specific reactivation refers to multivoxel reactivation of neural activity that occurred during perception of specific images, whereas feature-specific reactivation refers to reactivation of neural activity that occurred during perception of specific features shared across images. Features were extracted from layer node activations using the VGG16 deep neural net (DNN) (Simonyan & Zisserman, 2014). Activations from the convolutional layers (1-13), and the fully connected layers (14-16) were used, corresponding to low-visual (edges and luminosity; 1-4), middle-visual (simple object parts and patterns; 5-9), high-visual (complex object parts, e.g. faces; 10-13) and semantic (object category; 14-16) features, respectively. The experiment (see Bone, Ahmad & Buchsbaum, 2020, which addressed a different question using the same experimental data) had two video viewing runs, used to train the encoding models, and three sets of alternating encoding and retrieval runs (Figure 1). During encoding runs participants memorized a set of thirty color images (per run) while performing a 1-back task. In the following retrieval runs, participants’ recall and recognition memory of the images was assessed. Neural reactivation was measured while participants visualized a cued image within a light-grey rectangle, followed by a memory vividness rating. An image was then presented that was either identical to the cued image or a similar lure, and the participants judged whether they had seen the image during encoding and provided a confidence rating. Critically, the lure images carried the semantic and visual gist of the cued images but had different fine-grained details (see Supplementary Figure 1 for example image pairs). Consequently, accuracy on the recognition task served as a measure of detailed, rather than gist-like, memory.

**Figure 1.**
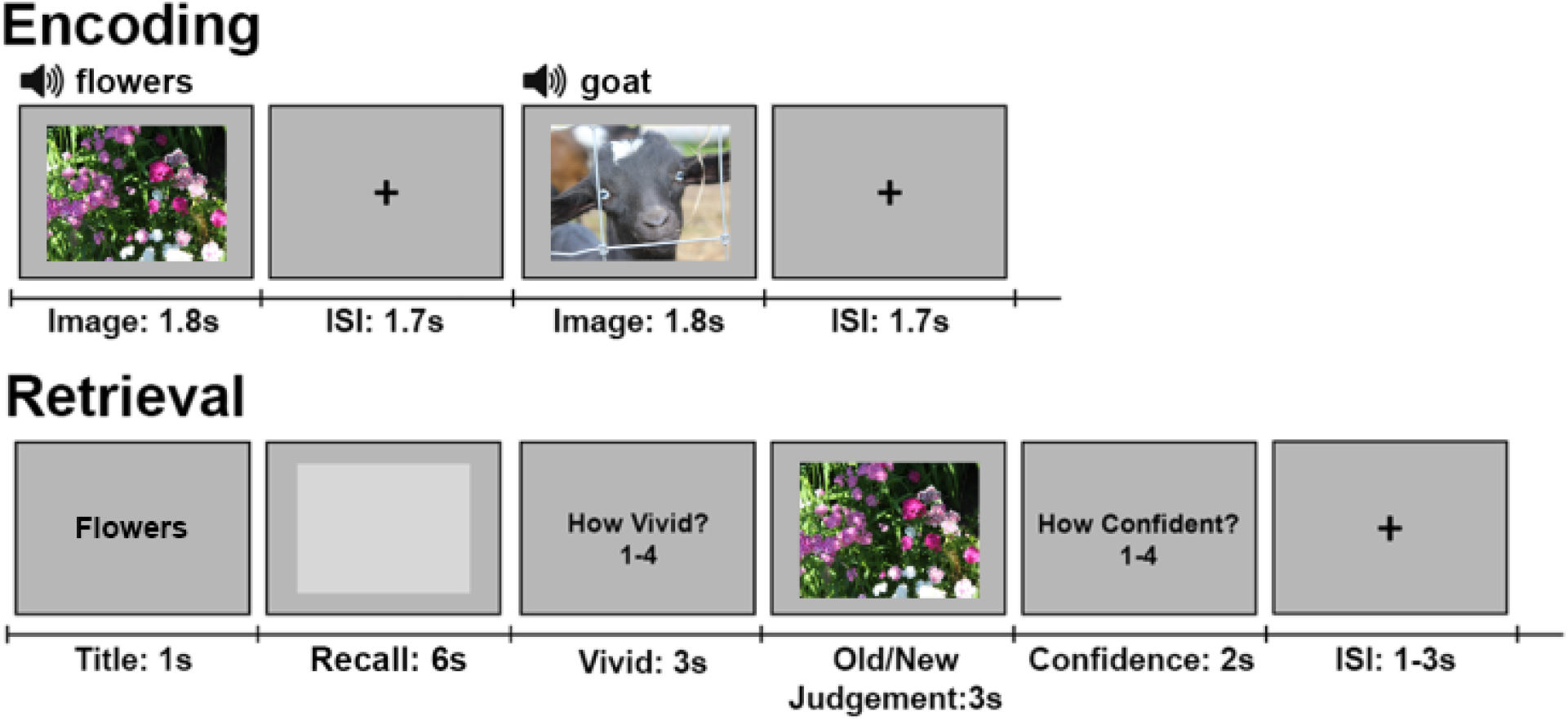
Procedure. Alternating image encoding and retrieval tasks. During encoding, participants performed a 1-back task while viewing a sequence of color photographs accompanied by matching auditory labels. During retrieval, participants 1) were cued with a visually-presented label, 2) retrieved and maintained a mental image of the associated photograph over a 6 second delay, 3) indicated the vividness of their mental image using 1-4 scale, 4) decided whether a probe image matched the cued image, and 5) entered their confidence rating with respect to the old/new judgement. Due to copyright concerns, images used in the study could not be included in the figure. The images depicted in the figure are for explanatory purposes only.

Hippocampal indexing theory claims that episodic memory retrieval involves hippocampal reactivation of an “index” that references a distributed neural representation stored in the neocortex. Assuming indexes with overlapping features are not completely ‘pattern separated’ (otherwise feature-specific reactivation would not occur within the hippocampus because feature-specific patterns would not be shared across events), episodic memory must require concurrent feature-specific reactivation within the hippocampus and regions of the neocortex that represent the target features. Recent experimental evidence (Bone, Ahmad & Buchsbaum, 2020) has shown that reactivation of low-level visual features within the early visual cortex is associated with greater visual recognition accuracy. Therefore, given the purported role of the pHC in representing detailed memories, we hypothesize that low-level visual reactivation within the pHC correlates with recognition accuracy only when accompanied by low-level visual reactivation within the early visual cortex (and vice-versa); i.e. recognition accuracy (and recall) is facilitated by the joint reactivation of low-level features within the hippocampus and neocortex.

We found that the within-subject correlation between recognition accuracy and image-specific reactivation during recall was significantly greater within the pHC relative to the aHC for individuals with high lure accuracy. Feature-specific reactivation during recall produced a similar result, with the correlations limited to low- and high-level visual features (e.g. edges and complex object parts). Moreover, low-level visual reactivation within the pHC positively interacted with low-level visual reactivation within the calcarine sulcus, indicating that the correlation between recognition accuracy and hippocampal reactivation depended upon reactivation co-occurring within the early visual cortex (and vice-versa). Overall, our results provide novel evidence for the indexing theory of hippocampal function and the role of the pHC in the representation of memory details. Our data also confirm and expand upon previous findings (Bone, Ahmad & Buchsbaum, 2020) showing large individual differences in the features underlying visual memory, and the impact of those differences on memory acuity.

## Materials and Methods

### Participants

Thirty-seven right-handed young adults with normal or corrected-to-normal vision and no history of neurological or psychiatric disease were recruited through the Baycrest subject pool, tested, and paid for their participation. Informed consent was obtained, and the experimental protocol was approved by the Rotman Research Institute’s Ethics Board. Subjects were either native or fluent English speakers and had no contraindications for MRI. Data from twelve of these participants was excluded from the final analyses for the following reasons: excessive head motion (5; removed if > 5mm within run maximum displacement in head motion), fell asleep (2), did not complete experiment (3), trial labeling error (1), second video run was cut short due to technical difficulties (1). Thus, twenty-five participants were included in the final analysis (13 males and 12 females, 20-32 years old).

### Stimuli

111 colored photographs (800 by 600) were gathered from online sources. For each image, an image pair was acquired using Google’s similar image search function, for a total of 111 image pairs (222 images). 21 image pairs were used for practice, and the remaining 90 were used during the in-scan encoding and retrieval tasks (see Supplementary Figure 1 for example image pairs). Each image was paired with a short descriptive audio title in a synthesized female voice (https://neospeech.com; voice: Kate) during encoding runs; this title served as a visually presented retrieval cue during the in-scan retrieval task. Two videos used for model training (720 by 480 pixels; 30 fps; 10m 25s and 10m 35s in length) comprised a series of short (∼4s) clips drawn from YouTube and Vimeo, containing a wide variety of themes (e.g. still photos of bugs, people performing manual tasks, animated text, etc.). One additional video cut from “Indiana Jones: Raiders of the Lost Ark” (1024 by 435 pixels; 10m 6s in length) was displayed while in the scanner, but the associated data was not used in this experiment because the aspect ratio (widescreen) did not match the images.

### Procedure

Before undergoing MRI, participants were trained on a practice version of the task incorporating 21 practice image pairs. Inside the MRI scanner, participants completed three video viewing runs and three encoding-retrieval sets. The order of the runs was as follows: first video viewing run (short clips 1), second video viewing run (short clips 2), third video viewing run (Indiana Jones clip), first encoding-retrieval set, second encoding-retrieval set, third encoding-retrieval set. A high-resolution structural scan was acquired between the second and third encoding-retrieval sets, providing a break.

Video viewing runs were 10m 57s long. For each run, participants were instructed to pay attention while the video (with audio) played within the center of the screen. The order of the videos was the same for all participants.

Encoding-retrieval sets were composed of one encoding run followed by one retrieval run. Each set required the participants to first memorize and then recall 30 images drawn from 30 image pairs. The image pairs within each set were selected randomly, with the constraint that no image pair could be used in more than one set. The image selected from each image pair to be presented during encoding was counterbalanced across subjects. This experimental procedure was designed to limit the concurrent memory load to 30 images for each of three consecutive pairs of encoding-retrieval runs.

Encoding runs were 6m 24s long. Each run started with 10s during which instructions were displayed on-screen. Trials began with the appearance of an image in the center of the screen (1.8s), accompanied by a simultaneous descriptive audio cue (e.g. a picture depicting toddlers would be coupled with the spoken word “toddlers”). Images occupied 800 by 600 pixels of a 1024 by 768 pixel screen. Between trials, a crosshair appeared centrally (font size = 50) for 1.7s. Participants were instructed to pay attention to each image and to encode as many details as possible so that they could visualize the images as precisely as possible during the imagery task. The participants also performed a 1-back task requiring the participants to press “1” if the displayed image was the same as the preceding image, and “2” otherwise. Within each run, stimuli for the 1-back task were randomly sampled with the following constraints: 1) each image was repeated exactly four times in the run (120 trials per run; 360 for the entire session), 2) there was only one immediate repetition per image, and 3) the other two repetitions were at least 4 items apart in the 1-back sequence.

Retrieval runs were 9m 32s long. Each run started with 10s during which instructions were displayed on-screen. Thirty images were then cued once each (the order was randomized), for a total of 30 trials per run (90 for the entire scan). Trials began with an image title appeared in the center of the screen for 1s (font = Courier New, font size = 30). After 1s, the title was replaced by an empty rectangular box shown in the center of the screen (6s), and whose edges corresponded to the edges of the stimulus images (800 by 600 pixels). Participants were instructed to visualize the image that corresponded to the title as accurately as they could within the confines of the box. Once the box disappeared, participants were prompted to rate the subjective vividness (defined as the relative number of recalled visual details specific to the cued image presented during encoding) of their mental image on a 1-4 scale (1 = a very small number of visual details were recalled, 4 = a very large number of visual details were recalled) (3s) using a four-button fiber optic response box (right hand; 1 = right index finger; 4 = right little finger). This was followed by the appearance of a probe image (800 by 600 pixels) in the center of the screen (3s), that was either the same as or similar to the trial’s cued image (i.e. either the image shown during encoding or its pair). While the image remained on the screen, the participants were instructed to respond with “1” if they thought that the image was the one seen during encoding (old), or “2” if the image was new (responses made using the response box). Following the disappearance of the image, participants were prompted to rate their confidence in their old/new response on a 1-4 scale (2s) using the response box. Between each trial, a crosshair (font size = 50) appeared in the center of the screen for either 1, 2 or 3 seconds.

Randomization sequences were generated such that both images within each image pair (image A and B) were presented equally often during the encoding runs across subjects. During retrieval runs each image appeared equally often as a matching (encode A −> probe A) or mismatching (encode A −> probe B) image across subjects. Due to the need to remove several subjects from the analyses, stimulus versions were approximately balanced over subjects.

### Setup and Data Acquisition

Participants were scanned with a 3.0-T Siemens MAGNETOM Trio MRI scanner using a 32-channel head coil system. Functional images were acquired using a multiband EPI sequence sensitive to BOLD contrast (22 x 22 cm field of view with a 110 x 110 matrix size, resulting in an in-plane resolution of 2 x 2 mm for each of 63 2-mm axial slices; repetition time = 1.77 sec; echo time = 30ms; flip angle = 62 degrees). A high-resolution whole-brain magnetization prepared rapid gradient echo (MP-RAGE) 3-D T1 weighted scan (160 slices of 1mm thickness, 19.2 x 25.6 cm field of view) was also acquired for anatomical localization.

The experiment was programmed with the E-Prime 2.0.10.353 software (Psychology Software Tools, Pittsburgh, PA). Visual stimuli were projected onto a screen behind the scanner made visible to the participant through a mirror mounted on the head coil.

### fMRI Preprocessing

Functional images were converted into NIFTI-1 format, motion-corrected and realigned to the average image of the first run with AFNI’s (Cox, 1996) 3dvolreg program. The maximum displacement for each EPI image relative to the reference image was recorded and assessed for head motion. The average EPI image was then co-registered to the high-resolution T1-weighted MP-RAGE structural using the AFNI program align_epi_anat.py (Saad et al., 2009).

The functional data for each experimental task (video viewing, 1-back encoding task, retrieval task) was then projected to a subject-specific cortical surface generated by Freesurfer 5.3 (Dale, Fischl & Sereno, 1999). The target surface was a spherically normalized mesh with 32000 vertices that was standardized using the resampling procedure implemented in the AFNI program MapIcosahedron (Argall, Saad & Beauchamp, 2006). To project volumetric imaging data to the cortical surface we used the AFNI program 3dVol2Surf with the “average” mapping algorithm, which approximates the value at each surface vertex as the average value among the set of voxels that intersect a line along the surface normal connecting the white matter and pial surfaces.

The three video scans (experimental runs 1-3), because they involved a continuous stimulation paradigm, were directly mapped to the surface without any pre-processing to the cortical surface. The three retrieval scans (runs 5, 7, 9) were first divided into a sequence of experimental trials with each trial beginning (t=-2) two seconds prior to the onset of the retrieval cue (verbal label) and ending 32 seconds later in two second increments. These trials were then concatenated in time to form a series of 90 trial-specific time-series, each of which consisted of 16 samples. The resulting trial-wise data blocks were then projected onto the cortical surface. To facilitate separate analyses of the “recall” and “old/new judgment” retrieval data, a regression approach was implemented. For each trial, the expected hemodynamic response associated with each task was generated by convolving a series of instantaneous impulses (i.e. a delta function) over the task period (10 per second; imagery: 61; old/new: 31) with the SPM canonical hemodynamic response. Estimates of beta coefficients for each trial and task were computed via a separate linear regression per trial (each with 16 samples: one per time point), with vertex activity as the dependent variable, and the expected hemodynamic response values for the “recall” and “old/new judgment” tasks as independent variables. The “recall” beta coefficients were used in all subsequent neural analyses. Data from the three encoding scans (runs 4, 6, 8) were first processed in volumetric space using a trial-wise regression approach, where the onset of each image stimulus was modelled with a separate regressor formed from a convolution of the instantaneous impulse with the SPM canonical hemodynamic response. Estimates of trial-wise beta coefficients were then computed using the “least squares sum” (Mumford, Turner, Ashby & Poldrack, 2012) regularized regression approach as implemented in the AFNI program 3dLSS. The 360 (30 unique images per run, 4 repetitions per run, 3 total runs) estimated beta coefficients were then projected onto the cortical surface with 3dVol2Surf.

### Hippocampal ROI definition

To define anterior and posterior hippocampal ROIs, we used the Freesurfer’s (version 5.3) automated parcellation of the left and right hippocampi on the T1-weighted image of each participant. To create anterior and posterior subsections of the hippocampus, these ROIs were equally divided into five sections along the antero-posterior axis, yielding five ROIs per hemisphere. These ROIs were then used to as masks to extract time-series from the pre-processed and co-registered fMRI data.

### Deep Neural Network Image Features

We used the pretrained TensorFlow implementation of the VGG16 deep neural network (DNN) model (Simonyan & Zisserman, 2014; see http://www.cs.toronto.edu/~frossard/post/vgg16 for the implementation used). Like AlexNet (the network used in previous studies, e.g. Güçlü & van Gerven, 2015), VGG16 uses Fukushima’s (1980) original visual-cortex inspired architecture, but with greatly improved top-5 (out of 1000) classification accuracy (AlexNet: 83%, VGG16: 93%). The network’s accuracy was particularly important for this study because we did not hand-select stimuli (images and video frames) that were correctly classified by the net. The VGG16 model consists of a total of thirteen convolutional layers and three fully connected layers. 90 image pairs from the memory task and 3775 video frames (3 frames per second; taken from the two short-clip videos; video 1: 1875 frames; video 2: 1900 frames; extracted using “Free Video to JPG Converter” https://www.dvdvideosoft.com/products/dvd/Free-Video-to-JPG-Converter.htm) were resized to 224 × 224 pixels to compute outputs of the VGG16 model for each image/frame. The outputs from the units in all layers were treated as vectors corresponding to low-level visual features (layers 1-4), mid-level visual features (layers 5-9), high-level visual features (layers 10-13) and semantic features (layers 14-16).

Convolutional layers (layers 1-13) were selected to represent visual features because they are modeled after the structure of the visual cortex (Fukushima, 1980), and previous work showed that the features contained within the convolutional layers of AlexNet (which has a similar architecture to VGG16) corresponded to the features represented throughout the visual cortex (Güçlü & van Gerven, 2015). The layer activations were visually inspected to confirm whether they represent the appropriate features. The low-level layers were required to have similar outputs to edge filters. Layers 1-4 best fit that description. The high-level layer was required to have features that selectively respond to complex objects (e.g. faces). Layers 10-13 contained such features. There were no a priori demands on the type of features represented by the middle layer, so layers 5-9 were selected. We used the fully connected layers (14-16) to approximate semantic features because, unlike the convolutional layers, they are not modeled upon the visual cortex. Instead, the fully connected layers are designed to learn features (derived from high-level visual features in layer 13) that directly contribute to the semantic classification of images.

To account for the low retinotopic spatial resolution resulting from participants eye movements, the spatial resolution of the convolutional layers (the fully connected layers have no explicit spatial representation) was reduced to 3 by 3 (original resolution for layers 1-2: 224 by 224; layers 3-4: 112 by 112; layers 5-7: 56 by 56; layers 8-10: 28 by 28; layers 11-13: 14 by 14). Convolutional layer activations were log-transformed to improve prediction accuracy (Naselaris et a., 2015).

### Feature-Specific Encoding Model

Separate encoding models were estimated for all combinations of subject, feature level and brain surface vertex (Naselaris et a., 2015). Let *v_it_* be the signal from vertex *i* during trial *t*. The encoding model for this vertex for a given feature level, *l*, is:

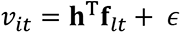

Here **f***_lt_* is a 100 × 1 vector of 100 image features from the layer of VGG16 representing the target feature level, *l*, associated with the current trial/image, *t* (only the 100 features from layer *l* with the largest positive correlations with the vertex activity, *v_i_*, were selected to make the computation tractable. Correlations were performed immediately before each non-negative lasso regression using data from the movie and encoding tasks), **h** is a 100 × 1 vector of model parameters that indicate the voxel/vertex’s sensitivity to a particular feature (the superscript T indicates transposition) and ε is zero-mean Gaussian additive noise.

The model parameters **h** were fit using non-negative lasso regression (R package “nnlasso”; Mandal & Ma, 2016) trained on data drawn from the encoding and movie viewing tasks (excluding the Indiana Jones video because its widescreen aspect ratio differed significantly from the encoded images) using 3-fold cross validation over the encoding data (cross validation was performed over images, so trials containing presentations of the to-be-predicted image were not included in the training set; all movie data was used in each fold). The non-negative constraint was included to reduce the possibility that a complex linear combination of low-level features may approximate one or more high-level features. The regularization parameter (lambda) was determined by testing 5 log-spaced values from approximately 1/10000 to 1 (using the nnlasso function’s path feature). For each value of the regularization parameter, the model parameters **h** were estimated for each vertex and then prediction accuracy (sum of squared errors; SSE) was measured on the held-out encoding data. For each vertex, the regularization parameter (lambda) that produced the highest prediction accuracy was retained for image decoding during recall.

### Image Decoding

For feature-specific reactivation, encoding models were used to predict neural activity during recall for each combination of subject, feature-level, ROI, and retrieval trial (148 cortical FreeSurfer ROIs and 10 hippocampal ROIs). The accuracy of this prediction was assessed as follows: 1) for each combination of subject, feature-level, and ROI the predicted neural activation patterns for the 90 images viewed during the encoding task were generated using a model that was trained on the movie and encoding task data, excluding data from encoding trials wherein the predicted image was viewed using 3-fold cross validation; 2) for each retrieval trial, the predictions were correlated (Pearson correlation across vertices within the given ROI) with the observed neural activity during recall resulting in 90 correlation coefficients. 3) the 90 correlation coefficients were ranked in descending order, and the rank of the prediction associated with the recalled image was recorded (1 = highest accuracy, 90 = lowest accuracy). 4) this rank was then subtracted from the mean rank (45.5) so that 0 was chance, and a positive value indicated greater-than-chance accuracy for the given trial (44.5 = highest accuracy, −44.5 = lowest accuracy). 5) the ranks were placed into four groups by layer (1-4, 5-9, 10-13, 14-16) and averaged together within each group, reducing the feature-levels from 16 to 4.

For image-specific reactivation, a similar decoding method was used (steps 1-4 ignoring references to feature-levels), except the predicted neural activation patterns for the 90 images were the average activation patterns (over four trials) when the participant viewed each image during encoding.

The reactivation results were averaged over bilateral ROI pairs (for cortical and hippocampal ROIs) to produce reactivation values for 74 bilateral cortical ROIs, and 5 bilateral hippocampal ROIs. To acquire anterior and posterior hippocampal reactivation values, the two most anterior and the two most posterior hippocampal ROIs (the middle ROI was not included) were averaged together, respectively.

### Removing Shared Variance Between ROIs and Feature Levels

To remove the shared variance between feature levels and the anterior and posterior hippocampus for the within-subject analyses, residuals extracted from linear models were used in the place of the reactivation measure. Linear models were run for all combinations of ROI and feature-level. For image-specific reactivation within the hippocampus, trial-by-trial reactivation within the aHC (pHC) was the DV and reactivation within the pHC (aHC) was the IV. For feature-specific reactivation within the neocortex, reactivation of the target feature level was the DV and reactivation of the three non-target feature levels were three IVs. For feature-specific reactivation within the hippocampus, reactivation of the target feature level within the aHC (pHC) was the DV, reactivation of the three non-target feature levels within the aHC (pHC) were three IVs, and reactivation of all feature levels within the pHC (aHC) were four IVs. The residuals from the models were used as measures of feature-specific reactivation in all within-subject analyses, replacing the reactivation measures used as the DVs.

### Bootstrap Statistics

For the within-subject LME models, confidence intervals and p-values were calculated with bootstrap statistical analyses (1000 samples) using the BootMer function (Bates, Maechler, Bolker & Walker, 2015). For the between-subject linear models, confidence intervals and p-values were generated with bootstrap statistical analyses (1000 samples) with random sampling over subjects.

### Data Availability

Data for all analyses covered in the article are available at https://github.com/MichaelBBone/ConcurrentReacHippoEV.

### Code Availability

Code for all analyses covered in the article are available at https://github.com/MichaelBBone/ConcurrentReacHippoEV.

## Results

### Recognition accuracy

Recognition accuracy, averaged across participants, was 83.3% (SD = 6.8%; chance = 50%). Accuracy on old and lure trials was 81.7% (SD = 9.8%) and 85.0% (SD = 10.7%), respectively, with no significant difference in accuracy between the two conditions (t(24) = −1.07, p = .295, paired samples, two-tailed t-test) (Supplementary Figure 2). Participants failed to respond within the three second old/new response period on 1.0% (SD = 1.5%) of trials. Those trials were classified as incorrect.

### Recognition accuracy and hippocampal reactivation during recall

We first set out to examine the relationship between trial-by-trial recognition accuracy and feature-specific reactivation within the anterior and posterior hippocampus, as well as whether this relationship differs between participants based upon their average lure accuracy. The authors previously found stark individual differences in the relation between recognition accuracy and low-level visual reactivation within the early visual cortex (Bone, Ahmad & Buchsbaum, 2020). Bone, Ahmad & Buchsbaum (2020) found that the subjects’ average recognition accuracy interacted with the trial-by-trial correlation between recognition accuracy and low-level visual reactivation within the early visual cortex such that the correlation was stronger for participants with greater accuracy. A follow-up analysis indicated that this interaction was specific to subjects’ average lure trial accuracy (no interaction was found for subjects’ average old trial accuracy). Consequently, we investigated the interaction between hippocampal reactivation and each subject’s average lure accuracy.

A binomial mixed linear effects (MLE) model was constructed with trial-by-trial accuracy as the dependent variable (DV), feature-specific reactivation of four feature levels (low-, mid-, high-level visual and semantic) within the aHC and pHC as eight independent variables (IV), the interaction between participants’ average lure accuracy and the four levels of feature-specific reactivation within the aHC and pHC as eight independent variables (IV), probe type (old or lure) as an IV (control), and subject and image pair as crossed random effects (random intercept only due to model complexity limitations). In addition to the feature-specific reactivation model, a similar binomial MLE model was constructed in which image-specific reactivation replaced the four levels of feature-specific reactivation. To focus on the unique contributions of the aHC, pHC, and the four feature levels, residuals extracted from linear models (with the variance shared between the aHC, pHC, and the four feature levels removed) were used in the place of the reactivation measures in the above analysis (see “Accounting for Shared Variance Between ROIs and Feature Levels” in Methods). Residuals were used in this way for all subsequent analysis. The coefficients from these models are depicted in Figure 2 (feature-specific: 2a-b, 2d-f; image-specific: 2c, 2g).

**Figure 2.**
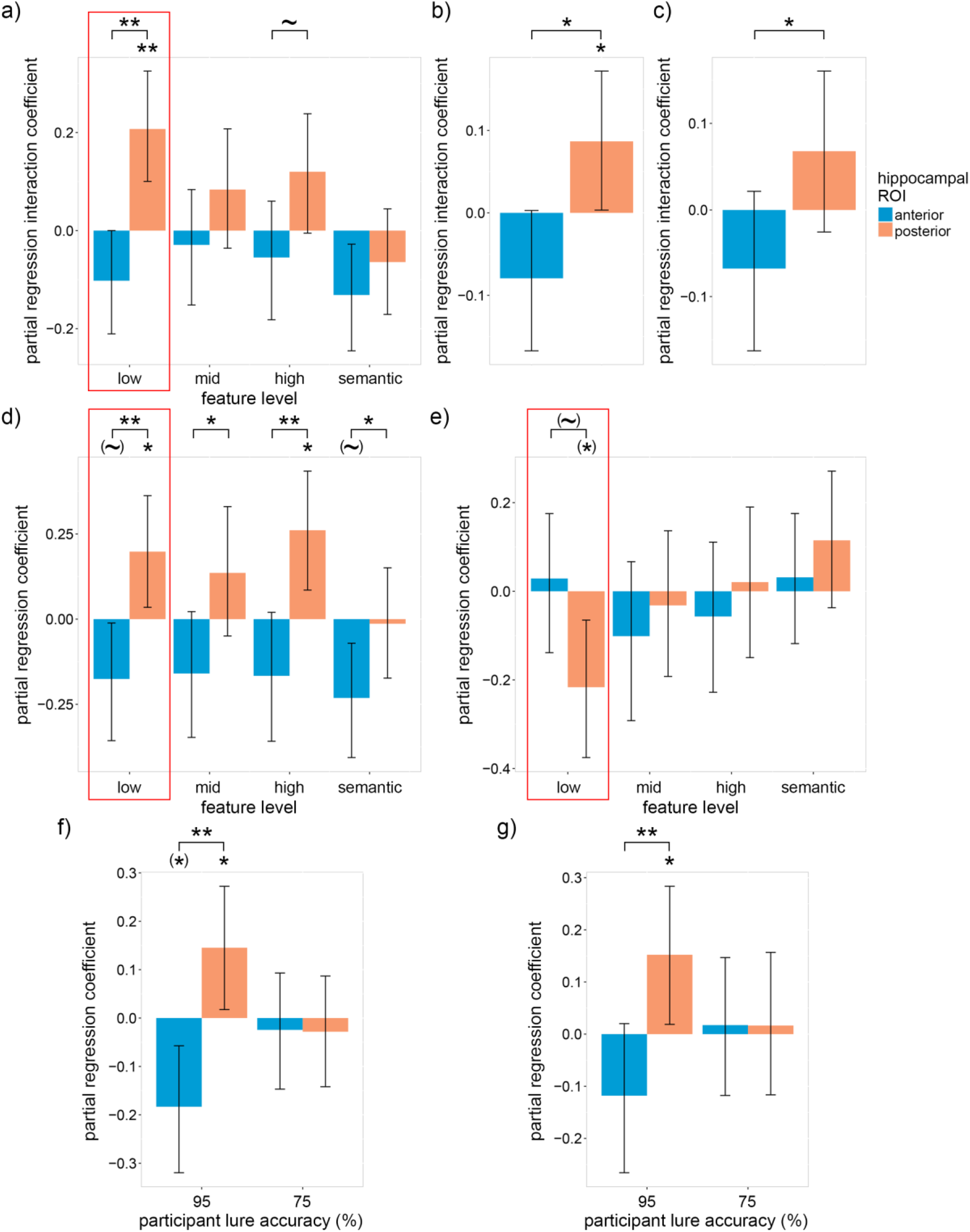
Relation between recognition accuracy and neural reactivation within the anterior and posterior hippocampus during recall. a) Within-subject partial regression coefficients for the interaction between feature-specific neural reactivation and each subjects’ average recognition lure accuracy with respect to trial-by-trial recognition accuracy. b) The coefficients in a) averaged over feature levels. c) The same interaction as a) except with item-specific (instead of feature-specific) reactivation. d) Within-subject partial regression coefficients from the same model as a) for the relation between feature-specific neural reactivation and trial-by-trial recognition accuracy for participant lure accuracy 1 standard deviation above average (95%). e) the same as d) but for participant lure accuracy 1 standard deviation below average (75%). f) The coefficients in d) and e) averaged over feature levels. g) The same coefficients as d) and e) except with item-specific (instead of feature-specific) reactivation. Error bars are 90% CIs; ∼ indicates p < 0.10, * indicates p < 0.05, ** p < 0.01, one-tailed bootstrap; (∼) indicates p < 0.10, (*) indicates p < 0.05, two-tailed bootstrap; FDR corrected over visual feature levels except for low-level features because, in accordance with our hypotheses, low-level features were prioritized (indicated by the red boxes).

Consistent with previous findings of individual differences within the visual cortex (Bone, Ahmad & Buchsbaum, 2020), a significant positive interaction was found between participants’ average lure accuracy and low-level visual reactivation within the pHC [β = .21, p = .001; one-tailed 1000 sample bootstrap—all bootstrap statistics used 1000 samples] (Figure 2a). In contrast, the interaction with low-level visual reactivation within the aHC was negative [albeit not significantly so: β = −.10, p = .102; two-tailed bootstrap] and significantly less than the pHC interaction [p = .001; paired-samples one-tailed bootstrap]. A qualitative analysis of the interaction coefficients in figure 2a indicates that this relationship between the pHC and aHC is consistent across feature levels. To assess this trend quantitatively, we averaged the coefficients across feature levels and found that the averaged pHC interaction coefficient was significantly greater than zero and the aHC coefficient [pHC: β = .09, p = .045; one-tailed bootstrap; aHC: β = −.08, p = .112; two-tailed bootstrap; difference: p = .010; paired-samples one-tailed bootstrap] (Figure 2b). Qualitatively similar albeit quantitatively weaker interaction coefficients were found using the image-specific reactivation model [pHC: β = .07, p = .120; one-tailed bootstrap; aHC: β = −.07, p = .204; two-tailed bootstrap; difference: p = .042; paired-samples one-tailed bootstrap] (Figure 2c). Overall, these findings reveal pronounced individual differences which suggest that reactivation within the posterior hippocampus, particularly low-level visual reactivation, is more strongly associated with recognition performance for participants with high lure accuracy.

To account for these striking individual differences, the partial regression coefficients for the relation between recognition accuracy and neural reactivation were assessed for participant lure accuracy one standard deviation above average (95%; Figure 2d) and one standard deviation below average (75%; Figure 2e). We first examine the results for individuals with high lure accuracy (Figure 2d). As hypothesized, the low-level visual reactivation coefficient within the pHC was significantly greater than zero and was also significantly greater than the corresponding aHC coefficient [pHC: β = .20, p = .021; one-tailed bootstrap; aHC: β = −.18, p = .076; two-tailed bootstrap; pHC - aHC: p = .006; paired-samples one-tailed bootstrap]. Looking beyond low-level features, we see that the pHC coefficient was significantly greater than zero for high-level features [β = .26, p = .018; one-tailed bootstrap; FDR corrected over feature levels], and the pHC coefficients were significantly greater than the aHC coefficients for all feature levels [mid-level visual: p = .022; high-level visual: p = .006; semantic: p = .045; paired-samples one-tailed bootstrap; FDR corrected over feature levels]. Qualitative analysis of Figure 2d suggests that interaction coefficients are generally positive within the pHC and negative within the aHC. The coefficients averaged over feature levels (Figure 2f) confirm this trend [pHC: β = .15, p = .031; one-tailed bootstrap; aHC: β = −.18, p = .024; two-tailed bootstrap; pHC - aHC: p = .004; paired-samples one-tailed bootstrap]. As with the interaction coefficients, the image-specific reactivation model produced qualitatively similar yet quantitatively weaker results [pHC: β = .15, p = .031; one-tailed bootstrap; aHC: β = −.12, p = .148; two-tailed bootstrap; pHC - aHC: p = .005; paired-samples one-tailed bootstrap] (Figure 2g). Taken as a whole, our results support the claim that the pHC represents memory details (which were expected to facilitate recognition accuracy), while the aHC represents more gist-like representations (which were expected to hinder recognition accuracy).

The above findings only held for individuals with high lure accuracy. For individuals with low lure accuracy (Figure 2e), we found the inversion of our hypothesis, i.e. the low-level visual coefficient within the pHC was significantly less than zero and marginally less than the aHC coefficient [pHC coefficient: β = −.22, p = .032; two-tailed bootstrap; aHC coefficient: β = .03, p = .380; one-tailed bootstrap; pHC - aHC difference: p = .060; paired-samples two-tailed bootstrap]. No other coefficients were significant. Our results suggest that reactivation of detailed low-visual information reduced recognition accuracy, forcing low-performing subjects to rely upon the reactivation of other (suboptimal) features. While this may seem paradoxical, reactivation of information can be misleading if that information is inaccurate, and the detailed/high-resolution information represented within the pHC would be more sensitive to inaccuracy than the gist-like/low-resolution information represented within the aHC.

If individuals with low lure accuracy are unable to accurately recall detailed low-level features within the pHC, how did they attempt to compensate? Although we see no positive coefficients within Figure 2e that does not necessarily mean that the participants did not recall the associated features. An alternative explanation for the null findings is that the features were not sufficient for the difficult recognition task, as we hypothesized for sematic features and the gist-like features of the aHC. If individuals with low lure accuracy relied to a greater extent upon these suboptimal features, then we should see a negative relationship between the participants’ average lure accuracy and feature-specific reactivation. To test this hypothesis, a between-subject linear model was constructed with participants’ average lure accuracy as the dependent variable (DV) and feature-specific reactivation of the four feature levels within the aHC and pHC as eight independent variables (IV).

As predicted, a significantly negative coefficient was found for low-level visual features within the aHC [β = −.08, p = .006; two-tailed bootstrap], whereas the coefficient for low-level features within the pHC was approximately zero [β = .01, p = .409; one-tailed bootstrap; pHC - aHC: p = .009; paired-samples one-tailed bootstrap] (Figure 3a). Positive coefficients were found for mid-level, high-level and semantic features within the pHC, indicating that individuals with high lure accuracy relied to a greater extent upon representations within the pHC [mid-level visual: β = .04, p = .025; high-level visual: β = .04, p = .025; semantic: β = .04, p = .025; one-tailed bootstrap; FDR corrected over feature levels]. The coefficients averaged over feature levels (Figure 3b) support this interpretation [pHC: β = .03, p = .024; one-tailed bootstrap; aHC: β = −.03, p = .340; two-tailed bootstrap; pHC - aHC: p = .037; paired-samples one-tailed bootstrap]. As with previous results, a variant of the above model using image-specific reactivation produced qualitatively similar yet quantitatively weaker results [pHC: β = .01, p = .347; one-tailed bootstrap; aHC: β = −.03, p = .150; two-tailed bootstrap; pHC - aHC: p = .075; paired-samples one-tailed bootstrap] (Figure 3c). Our findings support our hypothesis that individuals with low lure accuracy attempted to compensate for inaccurate low-level representations within the pHC by relying upon gist-like representations within the aHC, although the lack of a significantly positive low-level aHC coefficient within Figure 2d indicates that this strategy was unsuccessful for the current task—as expected.

**Figure 3.**
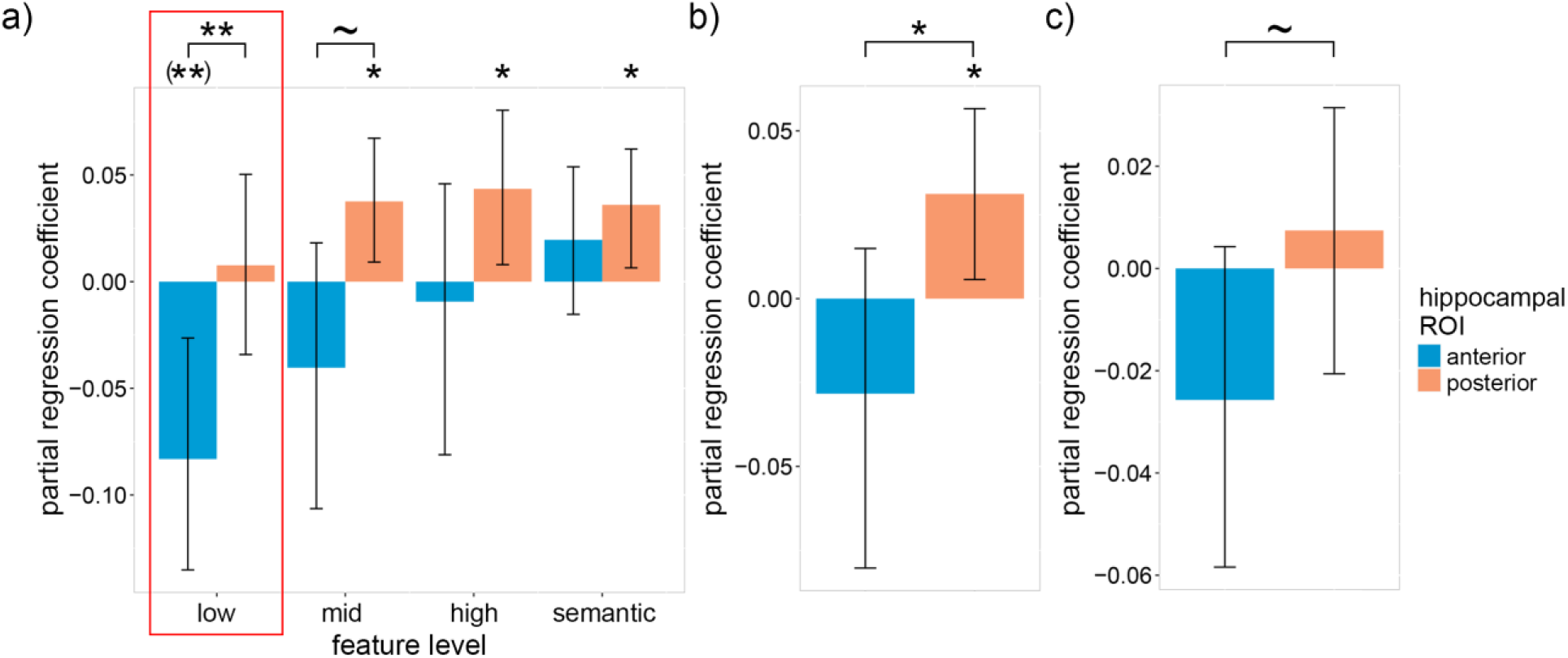
Relation between participants’ average recognition lure accuracy and neural reactivation within the hippocampus. a) Between-subject partial regression coefficients for the relation between feature-specific neural reactivation within the anterior and posterior hippocampus and each subjects’ average recognition lure accuracy. b) The coefficients in a) averaged over feature levels. c) The same coefficients as a) except with item-specific (instead of feature-specific) reactivation. Error bars are 90% CIs; ∼ indicates p < 0.10, * indicates p < 0.05, ** p < 0.01, one-tailed bootstrap; (**) indicates p < 0.01, two-tailed bootstrap; FDR corrected over visual feature levels except for low-level features because, in accordance with our hypotheses, low-level features were prioritized (indicated by the red boxes).

### Interaction between the hippocampus and calcarine sulcus

According to the hippocampal indexing theory (Teyler & DiScenna, 1986), episodic memories are encoded in distributed neural networks comprising hippocampal and neocortical neurons. Therefore, hippocampal reactivation should facilitate episodic recognition accuracy only when it co-occurs within relevant neocortical regions (and vice versa). To test this claim, a linear model was used to investigate whether a positive interaction exists between low-level visual reactivation within the pHC and calcarine sulcus (the neocortical region wherein V1 is concentrated; DeYoe et al. 1996), with respect to recognition accuracy (Figure 4a). We focused on low-level features for this analysis because low-level reactivation correlated with recognition accuracy (indicating that the features are useful for the task) and it is known a priori where low-level visual features are represented within the neocortex. Given that participants with low lure accuracy showed no relationship between recognition accuracy and hippocampal reactivation (Figure 2e-g), the interaction between the hippocampus and calcarine sulcus was expected only for participants with high lure accuracy.

**Figure 4.**
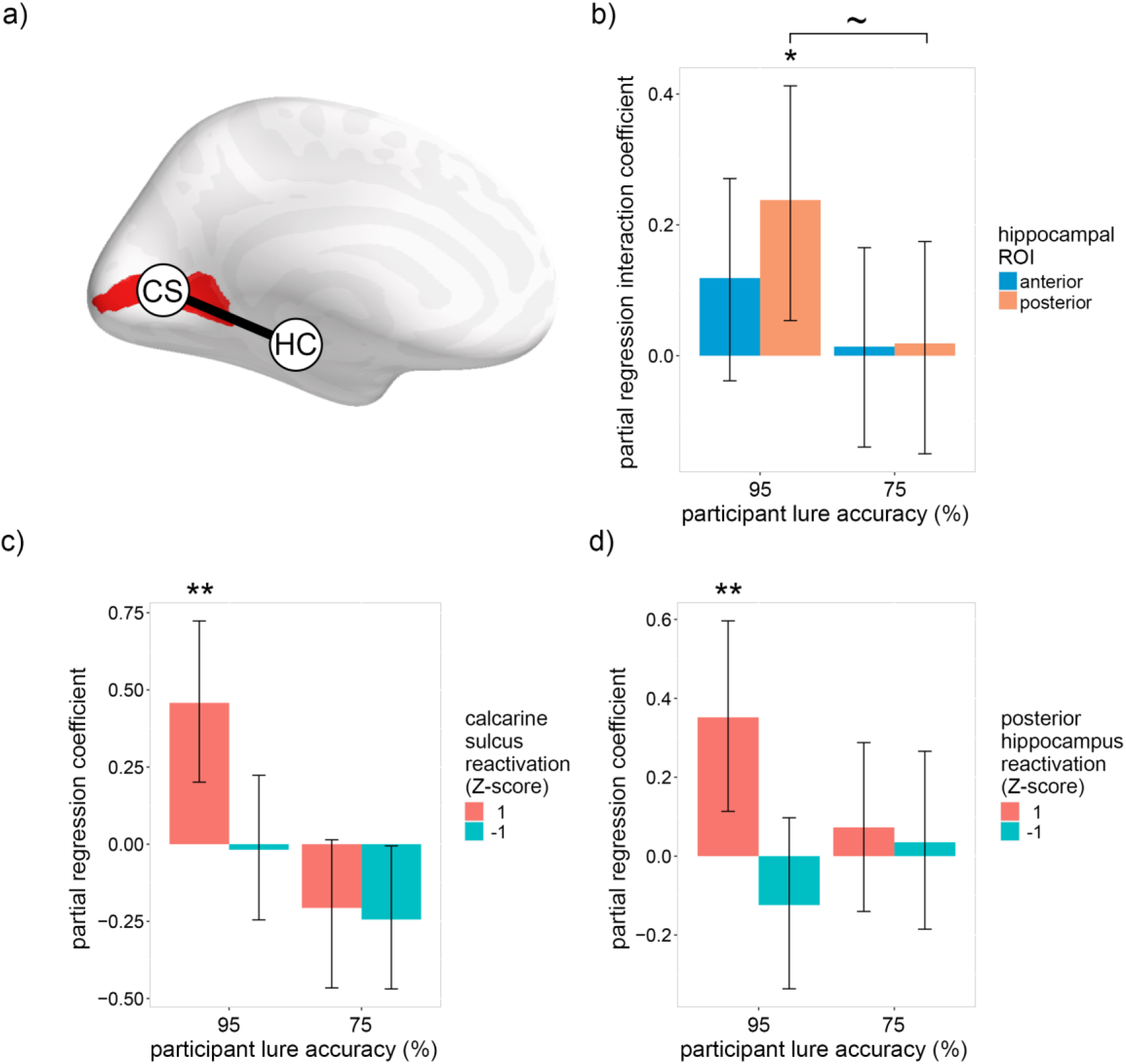
Relation between recognition accuracy and low-level visual neural reactivation within the hippocampus, calcarine sulcus and their interaction during recall. a) Diagram of the regions of interest: CS = calcarine sulcus, HC = hippocampus. The bilateral calcarine sulcus ROI is indicated in red. The line between the two ROIs represents the interaction between the regions. b) Within-subject partial regression coefficients for the interaction of low-level visual reactivation within the hippocampus and calcarine sulcus with respect to recognition accuracy. Interaction coefficients are displayed for participant lure accuracy 1 standard deviation above (95%) and below (75%) average. c) Within-subject partial regression coefficients from the same model as a) for the relation between low-level visual reactivation within the posterior hippocampus and recognition accuracy assuming participant lure accuracy 1 standard deviation above (95%) and below (75%) average and assuming low-level visual reactivation within the calcarine sulcus 1 standard deviation above and below average. c) Within-subject partial regression coefficients from the same model as a) for the relation between low-level visual reactivation within the calcarine sulcus and trial-by-trial recognition accuracy assuming participant lure accuracy 1 standard deviation above (95%) and below (75%) average and assuming low-level visual reactivation within the posterior hippocampus 1 standard deviation above and below average.

The binomial MLE model for the interaction between the hippocampus and calcarine sulcus consisted of trial-by-trial accuracy as the dependent variable (DV), reactivation of all four feature levels within the anterior and posterior hippocampus as 8 IVs, low-visual reactivation within the calcarine sulcus as an IV, the interactions between low-visual reactivation within the calcarine sulcus and all 8 hippocampal reactivation measures as 8 IVs, the interactions between participants’ average lure accuracy and all measures of reactivation (low-level within the calcarine sulcus and all levels within the aHC and pHC) as 9 IVs, the three-way interactions between average lure accuracy, low-visual reactivation within the calcarine sulcus and all 8 hippocampal reactivation measures as 8 IVs, probe type (old or lure) as an IV (control), and subject and image pair as crossed random effects (random intercept only due to model complexity limitations).

Figure 4b depicts the interaction between low-level visual reactivation within the calcarine sulcus and the hippocampus (see Supplementary Figure 3 for all interaction coefficients). As predicted by the hippocampal indexing theory, the pHC interaction coefficient was significantly greater than zero for individuals with high lure accuracy [pHC: β = .24, p = .017; aHC: β = .12, p = .105; one-tailed bootstrap]. For individuals with low lure accuracy, the interaction coefficients were approximately zero [pHC: β = .02, p = .466; aHC: β = .01, p = .476; one-tailed bootstrap], with the pHC coefficient marginally lower than the corresponding coefficient for individuals with high lure accuracy [p = .058; paired-samples one-tailed bootstrap]. To elaborate upon the observed pHC interaction, Figure 4c depicts the pHC partial regression coefficients when reactivation within the calcarine sulcus is either high (1; Z-scored) or low (−1), and Figure 4d depicts the calcarine sulcus partial regression coefficients when reactivation within the pHC is either high (1) or low (−1). For individuals with high lure accuracy, low-level visual reactivation within both hippocampal and early visual ROIs was positively associated with recognition accuracy only when reactivation within the other ROI was high [high reactivation in the other ROI: pHC: β = .46, p = .001; calcarine sulcus: β = .35, p = .007; one-tailed bootstrap; low reactivation in the other ROI: pHC: β = −.02, p = .890; calcarine sulcus: β = −.12, p = .352; two-tailed bootstrap]. The results indicate that trial-by-trial recognition accuracy was only associated with low-level visual reactivation when it co-occurred within the pHC and calcarine sulcus, thereby supporting the hippocampal indexing theory.

## Discussion

We investigated the relationship between performance on a recognition task that required retrieval of image-specific visual details and feature-specific reactivation within the hippocampus and neocortex. We showed that image-specific and feature-specific reactivation within the pHC, and not the aHC, positively correlated with recognition accuracy, indicating that the pHC indexes more detailed features relative to the aHC. Moreover, striking individual differences were observed such that recognition accuracy was positively associated with low-level visual reactivation within the pHC for individuals with above-average recognition lure accuracy, whereas the opposite relationship was observed for individuals with below-average recognition lure accuracy—which we postulate is caused by detailed low-level visual memories (e.g. the spatial location and orientation of edge features) being inaccurate to the point of being misleading in the latter group. Our results suggest that individuals with below-average recognition performance were not able to reactivate accurate low-level visual features within the pHC (i.e. representations with small receptive fields) and instead relied—ineffectually—upon the reactivation of less detailed low-level visual features within the aHC (i.e. representations with large receptive fields) which are more likely to overlap with the features of the lure images. Lastly, the correlation between recognition accuracy and low-level reactivation within the hippocampus was found to depend upon low-level reactivation within the early visual cortex (calcarine sulcus), and vice versa. This mutual dependence between hippocampal and neocortical feature-specific reactivation supports the hippocampal indexing theory.

The fact that it was possible to detect reactivation within the hippocampus using a model trained to recognize feature-specific patterns across images strongly implies that hippocampal indexes of different events are not entirely orthogonalized/pattern-separated (i.e. there must be some overlap between the indexes of memories that share similar features), but a couple of caveats must be considered. First, feature-specific reactivation could potentially be detected if orthogonal/pattern-separated indexes of related events (i.e. events with overlapping features, including, but not limited to, perception of the images in the current study) are consistently coactivated during encoding and retrieval. This interpretation does not align with our results because coactivation of indexes of similar events/images would be expected to reduce recognition accuracy, rather than facilitate it. Second, it may be the case that feature-specific reactivation was detected within hippocampal subregions that contain non-orthogonal representations. This interpretation is supported by studies (Bakker, Kirwan, Miller & Stark, 2008; Rolls, 2013) which suggest that orthogonal/pattern-separated representations are limited to the CA3 and dentate gyrus, whereas representations within the CA1 and subiculum are relatively non-orthogonal. Future work exploring feature-specific reactivation within individual hippocampal subregions will be required clarify how index orthogonality varies between subregions.

Multiple analyses, including image-specific and feature-specific reactivation both within- and between-subject, provided clear and consistent evidence that fine-grained visual features are represented within the pHC, whereas less detailed features are represented within the aHC. Although our results support and expand upon the majority of previous research using human and animal models (Kjelstrup et al., 2008; Schlichting, Mumford, & Preston, 2015; Evensmoen et al. 2015; Brunec et al., 2018; Sekeres, Winocur & Moscovitch, 2018; Grady, 2019), not all findings in the literature are consistent with this dichotomy. In a recent longitudinal study using a recognition task paradigm similar to the current study’s (except testing occurred 1 day or 28 days after encoding, and novel non-lure images were included in the recognition test), Dandolo and Schwabe (2018) found that detailed visual memories were represented within the aHC, whereas gist-like memories were represented within the pHC. The authors’ conclusion was based upon two findings: a between-subject correlation between lure accuracy and univariate activity within the hippocampus, and a multivariate model comparing cross-image neural pattern similarity for old, lure and new images. For the between-subject univariate correlation, recognition accuracy on lure trials positively correlated with univariate activity within the aHC, but not the pHC. In contrast, we found that participants’ average recognition accuracy on lure trials negatively correlated with reactivation within the aHC, and positively correlated with reactivation within the pHC. One potential explanation for this discrepancy is that memories may become more detailed within the aHC, relative to the pHC, over time (1 day). However, other work (Brunec et al., 2018) has found that representations within the aHC are less distinct than those within the pHC for memories encoded over periods much longer than 1 day. An alternative explanation is that univariate activity within the aHC during recognition may not reflect recollection of image-specific information, and instead reflects other functions such as context retrieval. The multivariate measure suffers from a similar issue of interpretation. Representational similarity analysis (RSA) was used to detect neural patterns shared between different images. Consequently, the detected effects do not represent retrieval of image-specific details, and instead might reflect the retrieval of the shared encoding context during recognition. Therefore, Dandolo and Schwabe’s (2018) univariate and multivariate results do not address whether the pHC or aHC represent task-relevant event-specific details, whereas our results based upon image/feature-specific reactivation do—and support the idea that the pHC and aHC represent detailed and gist-like memories, respectively.

Clear individual differences were observed with respect to the visual feature levels recalled and the detail/location of those features. For the recollection task, results suggest that individuals with high lure accuracy relied upon detailed low- and high-level visual reactivation within the pHC, and were hindered by semantic reactivation within the aHC, whereas low lure accuracy subjects ineffectually relied upon gist-like low-level reactivation within the aHC in an attempt to compensate for inaccurate low-level reactivation within the pHC. Our findings are consistent with and expand upon previous work by Sheldon et al. (2016) who found that people who tend to rely upon episodic memory (i.e. memories containing spatiotemporal and contextual details) have greater functional connectivity between the medial temporal lobes and posterior visual regions (i.e. neocortical regions predominately connected to the pHC; Poppenk, Evensmoen, Moscovitch, & Nadel, 2013). In contrast, people who tend to rely upon semantic memory were found to have greater functional connectivity between the medial temporal lobes and the prefrontal cortex (i.e. neocortical regions predominately connected to the aHC; Poppenk, Evensmoen, Moscovitch, & Nadel, 2013). Our findings are also consistent with Armson et al. (2019), who found that the association between eye fixation rate during free recall and the number of episodic details recalled was significantly greater for participants who tend to rely upon episodic memory (trait episodic and semantic memory in both studies was measured via questionnaire; Palombo, Williams, Abdi & Levine, 2013). It is reasonable to assume that the tendency to rely upon episodic or semantic memories would have an impact on the current study’s recognition task such that those who relied upon semantic memory would have relatively poor lure accuracy due to the strong semantic similarity between the encoded and lure images. It is therefore plausible that the individual differences observed within our study were driven by stable trait-like preferences for episodic or semantic memory. If this is the case, the current study suggests that those who favor semantic memory may do so because they are often unable to accurately recall detailed low-level perceptual features—possibly as a result of limited/impaired communication between the pHC and early visual cortex.

According to the hippocampal indexing theory, communication between the hippocampus and neocortex is necessary for episodic retrieval, at least before consolidation (Teyler & DiScenna, 1986). The theory posits that events are represented by unique spatiotemporal arrays of neocortical modules, and that these arrays are mapped/indexed by the hippocampus at encoding. During retrieval, the partial reactivation of an event’s neocortical array reactivates the hippocampal index associated with the event, which in turn reactivates the associated neocortical array, thereby simulating the original experience. Joint reactivation within the hippocampus and neocortex is therefore required for successful episodic retrieval. In support of the indexing theory, we found that for individuals with above-average lure accuracy the positive correlation between trial-by-trial recognition accuracy and low-level visual reactivation within the pHC depended upon low-level visual reactivation within the early visual cortex (calcarine sulcus), and vice-versa. Moreover, because our measure of feature-specific reactivation requires that similar features be associated with similar activity patterns across events, this suggests that the hippocampus contains a functionally topographic mapping of the neocortex (Teyler and DiScenna, 1986). Future reactivation studies should explore whether the observed hippocampal-neocortical interaction is evident for different features in the context of different tasks (the current task favored low-level visual features).

Individual differences provide additional support for the indexing theory. Participants with low lure accuracy had approximately equivalent low-level visual reactivation within the pHC relative to participants with high lure accuracy but showed no interaction between low-level visual reactivation within the hippocampus and calcarine sulcus. These findings suggest that individual differences in recognition memory may be the consequence of a functional disconnection between the hippocampus and neocortex. Future work should investigate why some individuals may lack the hippocampal-neocortical functional connections that appear to be necessary for detailed and accurate episodic memory.

To explain our findings more fully, we outline an expanded variant of the hippocampal indexing theory: Prediction Error Indexing (PEI). In addition to the indexing theory (Teyler & DiScenna, 1986), PEI incorporates ideas drawn from the complementary learning systems (CLS) theory (Kumaran, Hassabis & McClelland, 2016), the predictive coding theory (Rao & Ballard, 1999; Bastos et al., 2012) and the predictive, interactive multiple memory systems (PIMMS) theory (Henson & Gagnepain, 2010). In short, PEI assumes that hippocampal memory representations optimize compression by taking advantage of the event-specific information that overlaps with the statistical information stored within the neocortex. This is accomplished by biasing the features that are stored within the hippocampus to those that are not predicted by the neocortex at encoding, with the rest of the features reconstructed at retrieval via neocortical inference.

According to the predictive coding account of perception, perceptual experience results from the reciprocal exchange of bottom-up and top-down signals throughout the cortical hierarchy. During perception, top-down connections convey predictions, which are compared against the perceptual input to generate an error signal. This signal is then propagated back up the hierarchy to update the predictions and enhance memory of the features that diverged from expectations (Axmacher et al., 2010; Henson & Gagnepain, 2010). PEI posits that the hippocampus facilitates episodic memory by storing an index of the features that were not predicted by the neocortex (i.e. the difference between the final top-down prediction and the bottom-up input) at all levels of the representational hierarchy (including low-level visual features). An index of high-level conceptual features at the top of the representational hierarchy is also stored because a top-down prediction cannot be made without a top-level representation. During episodic memory recall, the hippocampal index of the higher-level features activate the corresponding neocortical representations which, in turn, are used to infer lower-level features, while the hippocampal index of the lower-level features serves to constrain this inference to be specific to the recalled event. Encoding and retrieval of higher-level features takes precedence over lower-level features because memory of lower-level features is usually dependent upon higher-level features (but not necessarily, e.g. when no higher-level feature is able to predict the lower-level features). Consequently, memory for lower-level features will show greater variability (within- and between-subject) because these features would tend to have the lowest utility relative to the space they take up in memory (the current task being an exception). Future work could test PEI by associating feature-specific hippocampal reactivation with the predictability of a stimuli’s features.

The contributions of this study were fourfold. First, we found that feature-specific representations within the hippocampus are functionally topographic and dependent upon feature-specific representations within the neocortex, thereby providing compelling evidence for the hippocampal indexing theory. Second, results from multiple analyses indicated that fine-grained and gist-like features are represented within the pHC and aHC, respectively. Third, individual differences in recognition lure accuracy were associated with striking differences in the relation between trial-by-trial memory accuracy and feature-specific reactivation within and between the hippocampus and neocortex. Fourth, we outlined a variant of the hippocampal indexing theory, Prediction Error Indexing (PEI), that is consistent with our results and incorporates recent theoretical advances in the fields of memory and perception. Overall, the current study’s results stress the importance of conceptualizing and measuring hippocampal function as part of an extended hippocampal-neocortical network, rather than in isolation. By doing so, future studies may refine our understanding of the mechanisms underlying memory and reveal the causes individual differences within healthy and clinical populations.

## End Notes

### Funding

This work was supported by the Natural Sciences and Engineering Research Council of Canada (488937 to B.R.B.) and the Canadian Institutes of Health Research (152879 to B.R.B).

## Acknowledgements

We thank Asaf Gilboa for his helpful input.

## Author Contributions

Conceptualization, M.B.B., B.R.B.; Methodology, M.B.B., B.R.B.; Software, M.B.B., B.R.B.; Formal Analysis, M.B.B.; Data Curation, M.B.B., B.R.B.; Writing – Original Draft, M.B.B.; Writing – Review and Editing, M.B.B., B.R.B.; Visualization, M.B.B.; Supervision, B.R.B.

## Competing Interests

The authors declare no competing interests.

## Notes

### Competing Interest Statement

The authors have declared no competing interest.

