## Supplemental Figure 1 for "Concurrent feature-specific reactivation within the hippocampus and neocortex facilitates episodic memory retrieval"

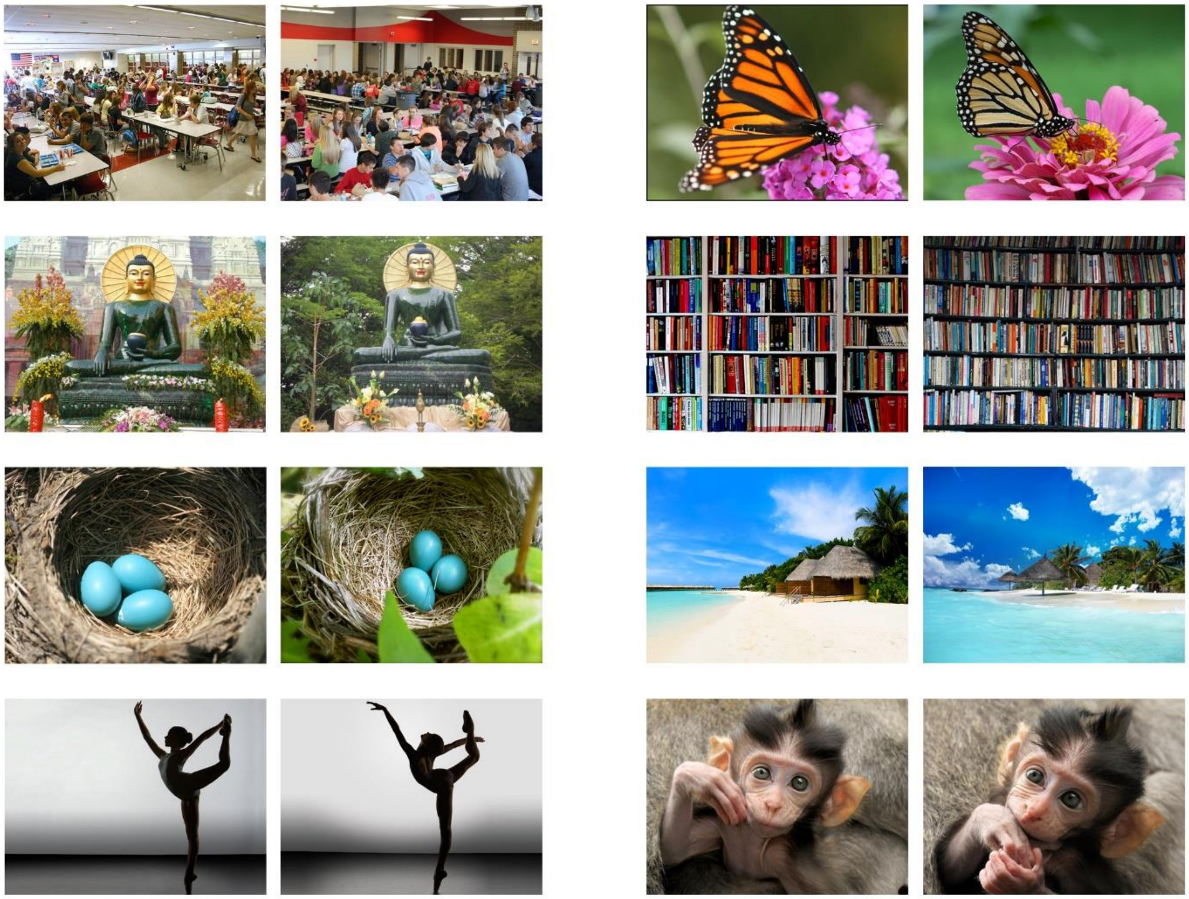

**Supplementary Figure 1. Example of Image Pairs.** Eight randomly selected image pairs out of the ninety total.
