## Supplementary figures and images for "Concurrent feature-specific reactivation within the hippocampus and neocortex facilitates episodic memory retrieval"

### Supplemental Figure 2

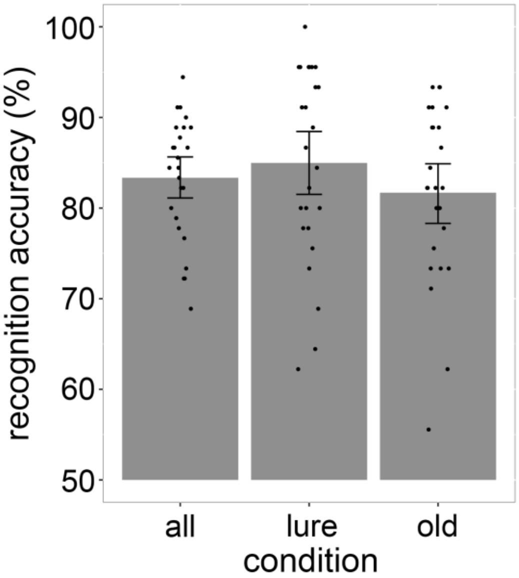


**Supplementary Figure 2. Recognition accuracy.** Each dot represents a subject. Error bars are 90% CIs.
