## Supplemental Figure 3 for "Concurrent feature-specific reactivation within the hippocampus and neocortex facilitates episodic memory retrieval"

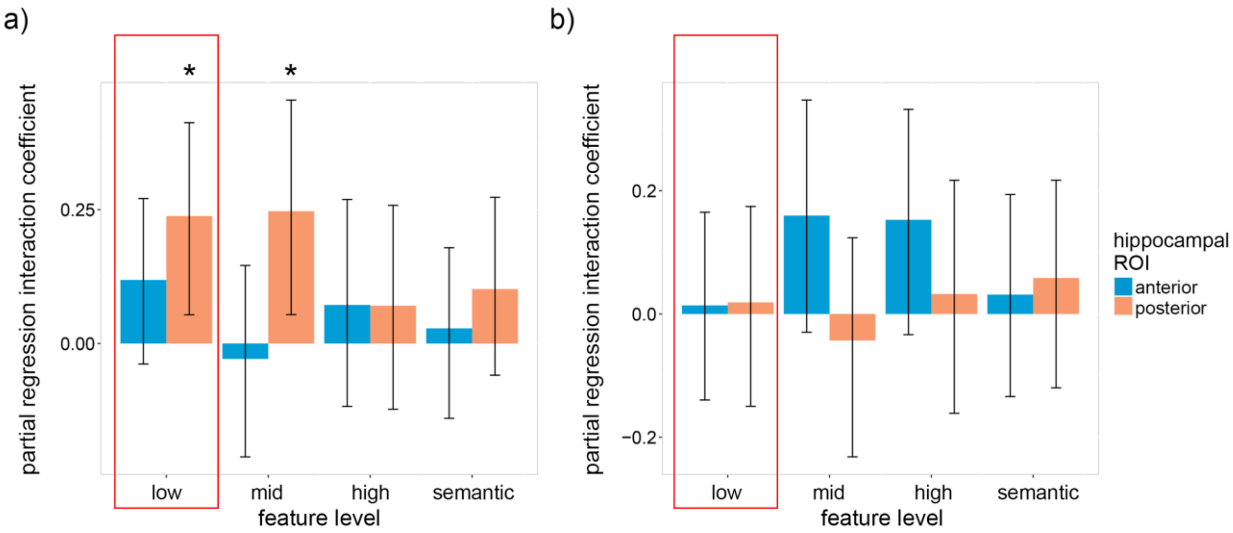


**Supplementary Figure 3. Interaction of feature-specific reactivation within the hippocampus and calcarine sulcus during recall with respect to recognition accuracy.** Within-subject partial regression coefficients for the interaction of feature-specific reactivation within the hippocampus and calcarine sulcus with respect to recognition accuracy given participant lure accuracy 1 standard deviation a) above (95%) and b) below (75%) average. The feature level along the x-axis refers to reactivation within the hippocampus. Reactivation within the calcarine sulcus was limited to low-level visual features. Error bars are 90% CIs; * indicates $p < 0.05$, one-tailed bootstrap; FDR corrected over visual feature levels except for low-level features because, in accordance with our hypotheses, low-level features were prioritized (indicated by the red boxes).
